# Antiviral activity of Glucosylceramide synthase inhibitors against SARS-CoV-2 and other RNA virus infections

**DOI:** 10.1101/2020.05.18.103283

**Authors:** Einat. B. Vitner, Roy Avraham, Hagit Achdout, Hadas Tamir, Avi Agami, Lilach Cherry, Yfat Yahalom-Ronen, Boaz Politi, Noam Erez, Sharon Melamed, Nir Paran, Tomer Israely

## Abstract

The need for antiviral drugs is real and relevant. Broad spectrum antiviral drugs have a particular advantage when dealing with rapid disease outbreaks, such as the current COVID-19 pandemic. Since viruses are completely dependent on internal cell mechanisms, they must cross cell membranes during their lifecycle, creating a dependence on processes involving membrane dynamics. Thus, in this study we examined whether the synthesis of glycosphingolipids, biologically active components of cell membranes, can serve as an antiviral therapeutic target. We examined the antiviral effect of two specific inhibitors of GlucosylCeramide synthase (GCS); (i) Genz-123346, an analogue of the FDA-approved drug Cerdelga®, (ii) GENZ-667161, an analogue of venglustat which is currently under phase III clinical trials. We found that both GCS inhibitors inhibit the replication of four different enveloped RNA viruses of different genus, organ-target and transmission route: (i) Neuroinvasive Sindbis virus (SVNI), (ii) West Nile virus (WNV), (iii) Influenza A virus, and (iv) SARS-CoV-2. Moreover, GCS inhibitors significantly increase the survival rate of SVNI-infected mice. Our data suggest that GCS inhibitors can potentially serve as a broad-spectrum antiviral therapy and should be further examined in preclinical and clinical trial. Analogues of the specific compounds tested have already been studied clinically, implying they can be fast-tracked for public use. With the current COVID-19 pandemic, this may be particularly relevant to SARS-CoV-2 infection.

**One Sentence Summary:** An analogue of Cerdelga®, an FDA-approved drug, is effective against a broad range of RNA-viruses including the newly emerging SARS-CoV-2.

## Introduction

Viral infections create a significant burden to human health worldwide, making the development of antiviral drugs a pressing need. Despite the rapid advancement in pharmaceutical and biotechnological approaches (e.g., RNA interference [RNAi] (*1*)), the development of successful antiviral treatments remains a challenge (*2*). Historically, drug research has mainly focused on targeting viral components, because of the perceived specificity of such an approach (*3*). However, since viral life cycle is dependent on the host, specific host mechanisms can also be explored as antiviral targets. There are distinct advantages for this approach such as creating a high barrier to resistance, providing broad coverage of different genotypes/serotypes, possibly even multiple viruses, and expanding the list of potential targets for a drug, when druggable viral targets are limited (*4*).

While side effects may be of particular concern for such treatments, another advantage of targeting host protein is the availability of many approved drugs against host proteins, allowing for drug repurposing. The main advantage of repurposing approved drugs is that they have already proven to be sufficiently safe, they have successfully passed clinical trials and regulatory scrutiny, and they have already undergone post-marketing surveillance (*5*).

This leads to significantly reduced timelines and required investment in making treatment available. In cases of major pandemic outbreaks caused by new viruses, shortening the time to treatment can have a major impact on public health and economics, making drug repurposing particularly desirable.

Sphingolipids (SLs) are biologically active components of cell membranes and as such are tightly linked to all processes involving membrane dynamics, making them potential key regulators in the life cycle of obligatory intracellular pathogens such as viruses. Glucosylceramide (GlcCer) is the backbone of more than 300 structurally different Glycosphingolipids (GSLs) including gangliosides and sulfatides. Its accumulation leads to Gaucher diseases accompanied by chronic brain inflammation and activation of the antiviral immune response (*6*). GSLs are involved in lateral and vertical segregation of receptors required for attachment, membrane fusion and endocytosis, as well as in intracellular replication, assembly and release of viruses. In addition, GSLs and their metabolites are inseparably interwoven in signal transduction processes, and the regulation of innate and intrinsic responses of infected target cells (*7*).

Viral-induced elevation of SL levels was shown to be associated with a number of viruses; elevation of GM2-ganglioside and Lactosylceramide was shown upon infection with Zika virus and Hepatitis C virus (HCV), respectively (*8, 9*). Human Cytomegalovirus (HCMV) induces elevation of ceramide and GM2-ganglioside (*10*), and Dengue virus induces elevation of ceramide and sphingomyelin (*11*). Additionally, Influenza virus was shown to induce Sphingomyelin and GlcCer elevation (*12, 13*) and suppression of the biosynthesis of cellular sphingolipids results in the inhibition of the maturation of influenza virus particles in vitro (*14, 15*). Moreover, iminosugars are known for their broad-spectrum antiviral activity, presumably because of their mechanism of action as endoplasmic reticulum (ER)-resident α-glucosidases I and II inhibitors (*16*). 1-Deoxynojirimycin (DNJ) iminosugar derivatives inhibit *in vitro* production of infectious viruses including dengue virus (DENV) (*17, 18*), hepatitis B virus (HBV) (*19, 20*), hepatitis C virus (HCV) (*21*), human immunodeficiency virus (HIV) (*22, 23*), and influenza A virus (*24*). Antiviral efficacy of the iminosugar *N*-butyl-DNJ (*N*B-DNJ, Miglustat, Zavesca) has been further demonstrated *in vivo* against DENV infection (*25*). Although these reports present strong circumstantial evidence that inhibition of ER α-glucosidase activity is the cause of iminosugar antiviral activity (*26*), the ubiquity of D-glucose in metabolism suggests that other pathways may be equally affected by iminosugar treatment. Indeed, *N*B-DNJ has been approved for clinical use since 2002 as a second line treatment for Gaucher’s disease (*27*) –a lysosomal storage disease (LSD). In this context, *N*B-DNJ is used as an inhibitor of UDP-glucose:ceramide glucosyltransferase (glucosylceramide synthase (GCS)), (EC 2.4.1.80) to reduce production of GSLs that accumulate due to a deficiency in GlcCer degradation (*28*). Thus, GSL synthetic pathways may be therapeutic targets for a broad-range of viral infection. Drugs targeting SL metabolizing enzymes are currently in use and constantly being developed for treating LSDs and other disorders in which alteration in SL levels are involved in disease pathology (31-29). This allows a potential repurposing of these already approved drugs as antivirals. While the inhibitor *N*B-DNJ affects multiple host targets, specific inhibition of GCS is now possible using GCS inhibitors which are currently available. In this study, we examined the antiviral activity of two specific inhibitors of GCS, which catalyze the biosynthesis of GlcCer. These inhibitors block the conversion of ceramide to GlcCer, the first step in the biosynthesis of gangliosides and other glycosphingolipids.

The following GCS inhibitors were examined: (i) (1R,2R)-nonanoic acid[2-(2′,3′-dihydro-benzo [1,4] dioxin-6′-yl)-2-hydroxy-1-pyrrolidin-1-ylmethyl-ethyl]-amide-l-tartaric acid salt (Genz-123346) termed herein after GZ-346. GZ-346 is an analogue of the FDA-approved drug eliglustat (Cerdelga®) which is indicated for the long-term treatment of adult patients with Gaucher disease type 1 (GD1) (*32*). (ii) (S)- quinuclidin-3-yl (2-(2-(4-fluorophenyl)thiazol-4-yl)propan-2-yl)carbamate (GENZ-667161) termed herein after GZ-161. GZ-161 is a specific inhibitor of glucosylceramide synthase (GCS) that can access the central nervous system (CNS) and has been demonstrated to effectively reduce glycosphingolipid synthesis (*33-35*). GZ-161 is an analogue of venglustat which is currently under clinical trials for the LSDs; Gaucher’s disease, Fabry disease, and Tay-Sachs disease, and is in a Phase 3 pivotal trial for autosomal-dominant polycystic kidney disease (*32-35*).

While initially our research was focused on virus-induced diseases of the CNS, the current focus on coronavirus disease 2019 (COVID-19) pandemic, have led us to test GCS inhibitors on non-neuronopathic viruses such as Influenza and severe acute respiratory syndrome coronavirus 2 (SARS-CoV-2), the virus causing COVID-19.

Thus, the antiviral activity of GCS inhibitors was examined on four enveloped RNA viruses, differing by their genus, organ-target and transmission route; (i) Neuroinvasive Sindbis virus (SVNI), a prototypic member of the alphavirus genus that has been used to study the pathogenesis of acute viral encephalitis in mice for many years (*36-38*). (ii) West Nile virus (WNV), a neurotropic flavivirus that has been the leading cause of arboviral encephalitis worldwide. (iii) Influenza A virus, a member of the family Orthomyxoviridae that can cause acute respiratory disease, and (iv) SARS-CoV-2, a members of the *Coronaviridae* family of the order Nidovirales (*39*). With the current coronavirus disease (COVID 19) outbreak, the potential effect of the tested compound is of particular interest and priority. Furthermore, Coronaviruses are behind two more large scale pandemic outbreaks in the past two decades: severe acute respiratory syndrome (SARS) in 2003, and the middle east respiratory syndrome (MERS) in 2012. The pandemic potential of these viruses along with the threat to public health they pose have put them on the WHO blueprint list for priority pathogens. The observed broad spectrum of effect of the tested compound makes it a good candidate for treatment of both current and future high priority pathogens.

## Results

### Antiviral activity of GCS inhibitors against SVNI virus

To determine whether GCS inhibitors block the replication of SVNI, Vero cells were incubated with serial dilutions of GZ-161 or GZ-346 one hour prior to infection with SVNI expressing luciferase (TRNSV-Luc) (*41*). Both GZ-161 and GZ-346 exhibited antiviral activity with an average median inhibitory concentration (IC50) of ∼4.5 μM and ∼7 μM, respectively (Fig.1A). No cytoxicity was observed in similarly treated uninfected cultures across the dose range (50% cytotoxic concentration (CC50) of >30 μM for both GZ-161 and GZ-346). Next, we examined the antiviral activity of GCS inhibitors on a mouse neural crest-derived cell line, Neuro 2A (N2a) to demonstrate that their antiviral activity is not specific to Vero cells. GZ-161 and GZ-346 inhibit SVNI replication in both cell lines, with percentage of inhibition of ∼40-50% and ∼60% in Vero and N2a cells, respectively (Fig 1B).

**Fig. 1.**
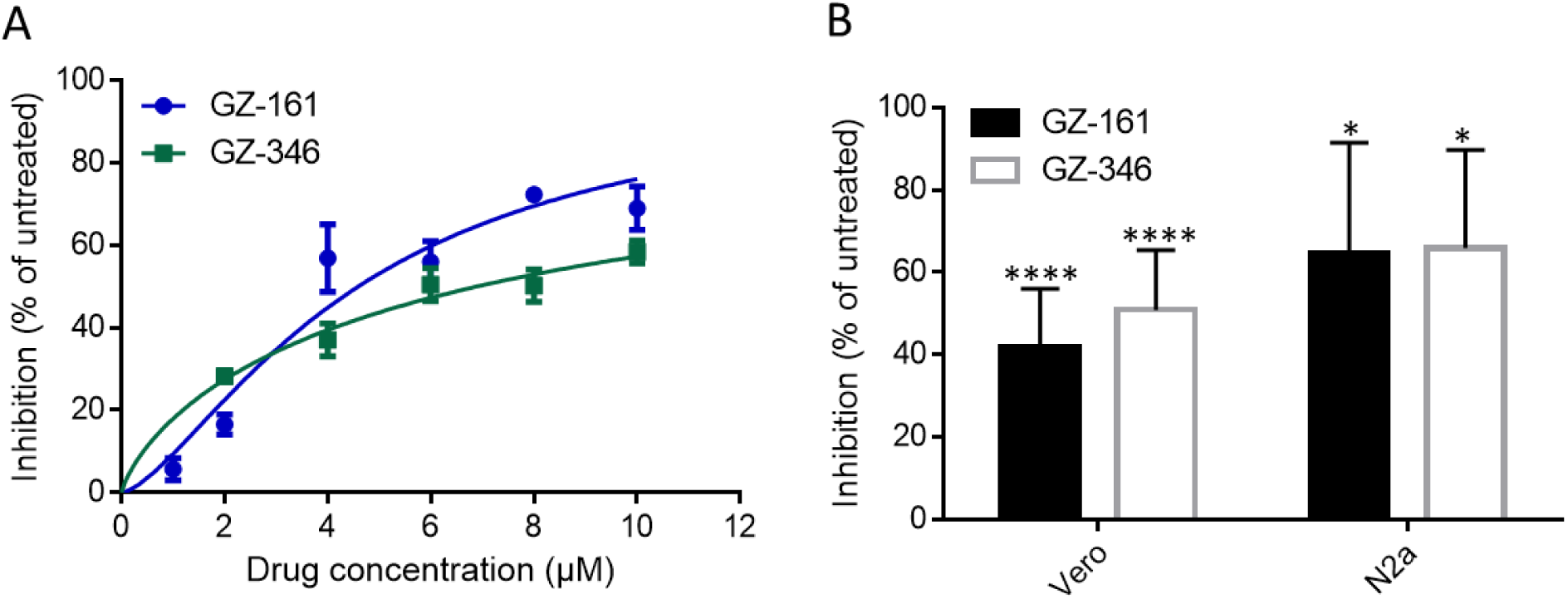
Inhibition of SVNI by GCS inhibitors *in-vitro*. (**A**) Dose-response curves of GZ-161 and GZ-346 for inhibiting recombinant neuroadapted Sindbis virus expressing Luciferase (TRNSV-Luc) infection. Vero cells were treated with GZ-161 or with GZ-346 (1-10µM). Cells were infected 1 hour later with TRNSV-Luc, MOI, 0.01. The infected cells were lysed 23 hours later, and the luciferase activities were measured. Data are means of four replicates ± SEM. I.C. 50 values are 4.5 µM and 6.9 µM for GZ-161 and GZ-346, respectively. (**B**) GCS inhibitors inhibit SVNI replication in both Vero and Neuro 2A (N2a) cells. Vero and N2a cells were seeded at a density of 3 × 10^4^ cells per well in 96-well plates. After incubating overnight, cells were treated with GZ-161 or with GZ-346 (10µM). Cells were infected 1 hour later with TRNSV-Luc, MOI, 0.01. The infected cells were lysed 23 hours later, and the luciferase activities were measured. Data are means of six replicates ± SEM. * p <0.05, **** p <0.0001 versus infected untreated.

### GCS inhibitors disrupt early stages of SVNI replication

To determine which stage of SVNI infection cycle was affected by GCS inhibitors, a time-of-addition assay was performed. As shown in Fig 2A, inhibition was most efficient when GCS inhibitors were added not later than 1 hour post infection (hpi) (Fig. 2A). The time-dependence of the inhibitory effect of the compound suggest that its anti-SVNI activity may be due to inhibition of early steps in the SVNI replication cycle. To further elaborate on the mechanism of GCS inhibition of SVNI life cycle, a Sindbis virus encoding green fluorescent protein (GFP) gene under the control of the subgenomic promoter (SIN-GFP) (*42*) was used. In this virus, GFP translation is only triggered upon the assembly of viral replication complexes enabling transcription of the sub-genomic RNA and translation of the encoded genes, including GFP in this case. To ensure single-cycle infection, a high multiplicity of infection (MOI) of 5 was applied. Vero cells were incubated with 10 μM GZ-161 or GZ-346 one hour prior to infection with SIN-GFP and GFP signal was measured at intervals of 2 hours (Fig. 2B). While GFP expression was highly elevated in untreated cells, a robust reduction was observed in GZ-161 and GZ-346 treated cells compared to untreated cells (Fig. 2B). To determine if GFP-levels were reduced due to a decrease in the number of GFP-expressing cells or due to an overall impaired expression in all cells, flow cytometry analysis of single-cycle infection was used (Fig. 2C). GZ-161 or GZ-346 significantly reduced the number GFP-expressing cells 24 hpi from ∼90% in untreated cell to ∼60% and ∼30% in GZ-161 and GZ-346 treated cell, respectively. Thus, our data suggest that GCS inhibitors inhibit SVNI infection cycle after attachment of the virus and before the translation of sub-genomic proteins.

**Fig. 2.**
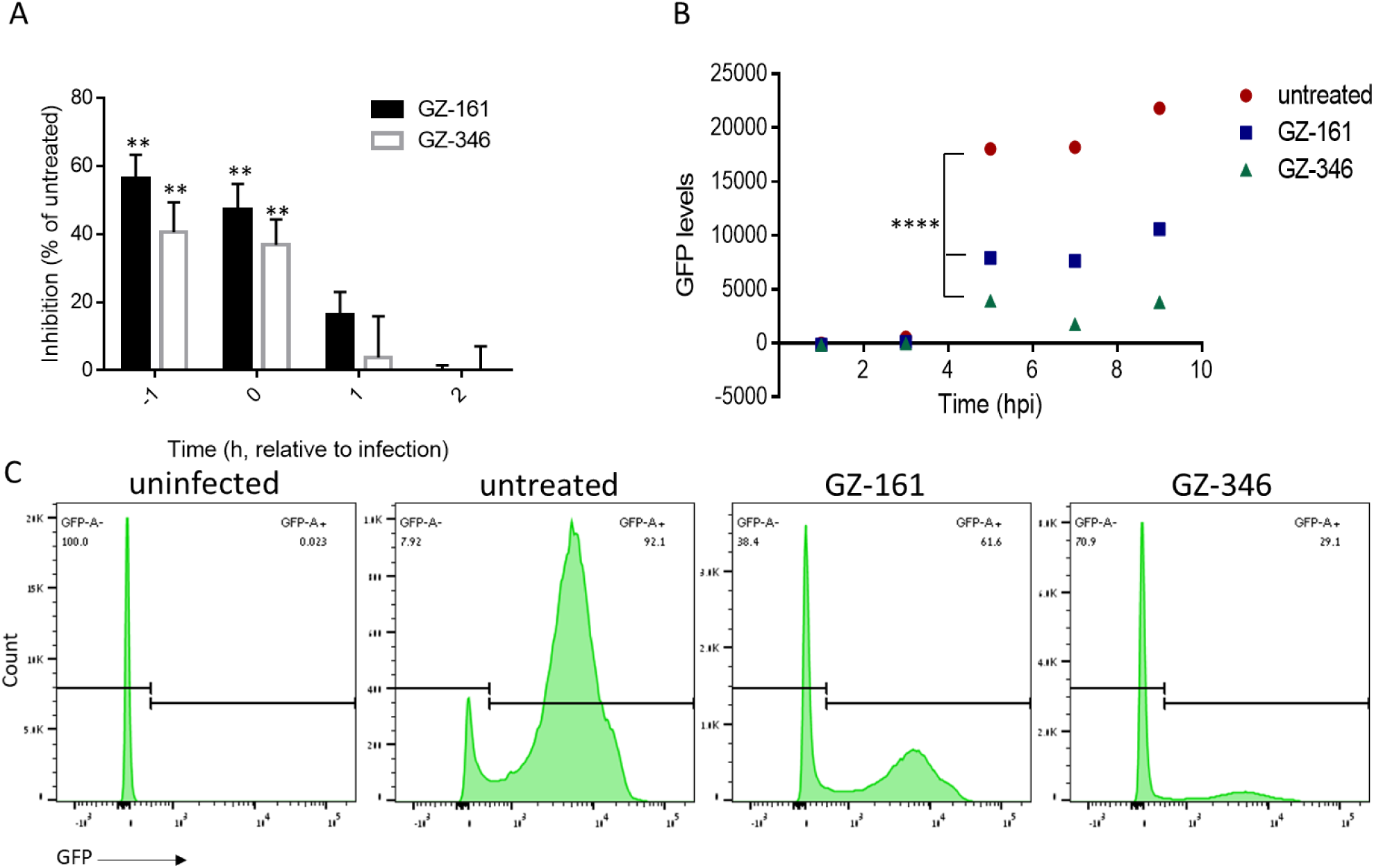
GCS inhibitors inhibit early stages of SVNI replication. **(A)** Time-of drug-addition assay. Vero cells were treated with 10 µM GZ-161 or GZ-346 1 hour prior to infection (−1), immediately post infection (0), 1 (1) or 2 (2) hours post infection. Cells were infected with TRNSV-Luc, MOI, 0.01, on ice for 1 hour following wash. The infected cells were lysed 23 hours later, and the luciferase activity was measured. Data are means of twelve replicates ± SEM. ** p <0.01 versus infected untreated. (**B**) Inhibition of subgenomic GFP expression by GCS inhibitors. Vero cells were treated with 10 µM GZ-161 or GZ-346 1 hour prior to infection. Cells were infected with SIN-GFP, MOI, 5, on ice for 1 hour following a wash. GFP levels were measured by microplate reader at the indicated time points post infection. Data are means of sixteen replicates ± SEM. **** p <0.0001 (**C**) Vero cells were treated with 10 µM GZ-161 or GZ-346 1 hour prior to infection. Cells were infected with SIN-GFP, MOI, 5, on ice for 1 hour following wash. 24 hpi cells were analyzed by flow cytometry. Data are representative of 3-replicates.

### GZ-161 enhances the survival of SVNI-infected mice

To evaluate whether GZ-161 could protect SVNI-infected mice, at a concentration comparable with the pre-clinical/clinical studies for drug approval, mice were treated with GZ-161 (20mg/kg/day, i.p) and infected with a lethal dose of SVNI. GZ-161 significantly protected pretreated infected mice from SVNI-induced mortality (Fig 3). Virtually all SVNI-infected mice died within 9 days post infection (∼90% mortality, with median time of survival (T50) of 7 days), whereas 60% of GZ-161-treated SVNI-infected animals survived (with undefined median survival) (Fig. 3). Postponing the treatment to the second day after infection still preserved some level of protection to SVNI-infected mice (Fig.3): 40% of the mice receiving post infection treatment with GZ-161 survived (with median survival of 9.5 days) (Fig. 3). Remarkably, both treatment regimens resulted in enhanced survival when compared to control, untreated group, with decreased mortality rates and increased median time of survival.

**Figure 3.**
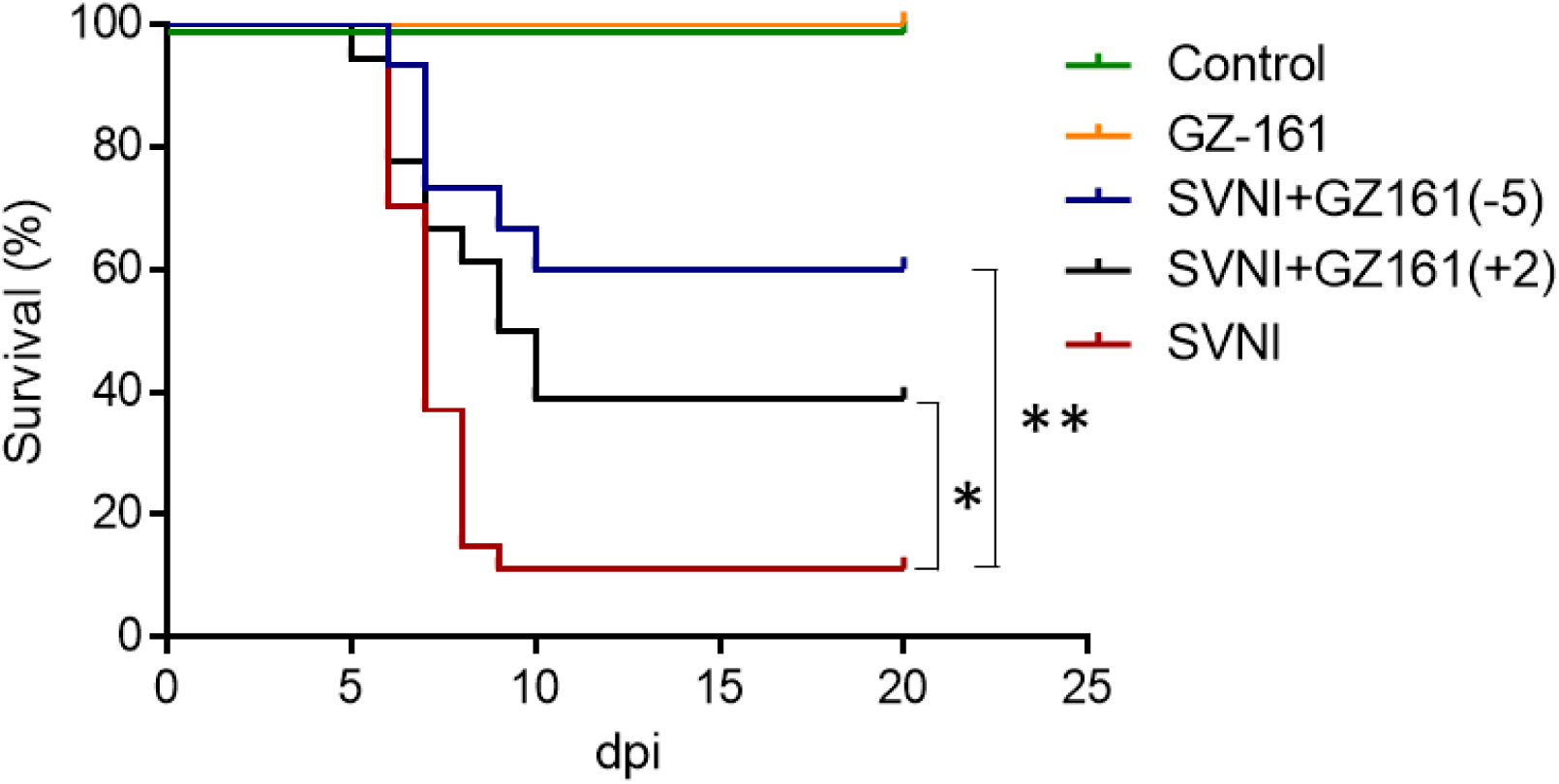
GZ-161 extends survival of SVNI-infected mice. (A) Kaplan–Meier survival curves of SVNI-infected mice (15 plaque forming units (PFU), administered intraperitoneally (i.p)). Mice were untreated (SVNI, n=27) or treated with GZ-161 (20 mg/kg/day, i.p), beginning at day 5 pre-infection (SVNI+GZ-161 (−5), n=15) or at day 2 post-infection (SVNI+GZ-161 (+2), n=18). Control mice were uninfected (n=6). Log-rank test for comparisons of Kaplan-Meier survival curves indicated a significant decrease in the mortality of GZ-161 treated mice compared to SVNI mice * p <0.05, ** p <0.01.

### Antiviral activity of GCS inhibitors against West nile virus

Next, we examined whether the antiviral activity of GCS inhibitors is specific to SVNI or whether they can block a neurotropic virus from different genus, namely WNV. Vero cells were incubated with 10 μM GZ-161 or GZ-346 one hour prior to infection with the NY-99 strain of WNV. Supernatants were harvested 24 hpi and analyzed by qPCR. We observed a 60% reduction in viral RNA present in the supernatant of samples treated with GZ-161 or GZ-346 compared to the vehicle DMSO (untreated), suggesting that GCS inhibitors has a broad antiviral activity towards neuronopathic RNA-viruses (Fig. 4).

**Figure 4.**
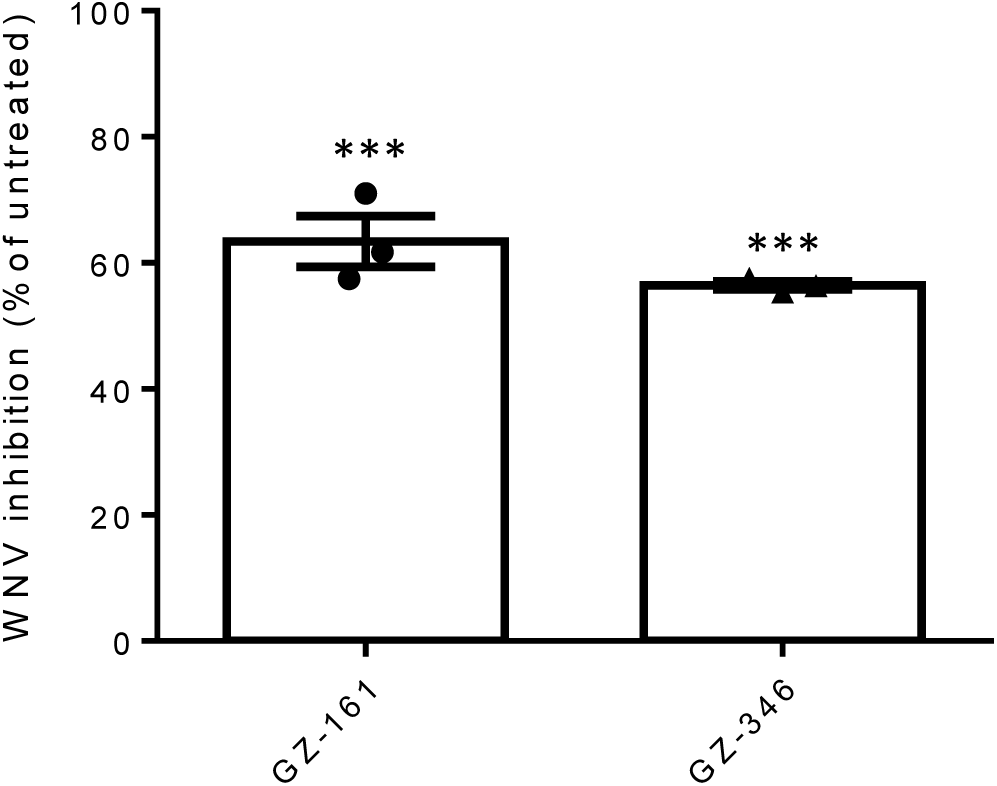
Inhibition of West Nile virus (WNV) by GCS inhibitors. Inhibition of WNV virus by glucosylceramide synthase inhibitors *in-vitro*. Vero cells were treated with GZ-161 or with GZ-346 (10µM). 1 hour later, cells were infected with WNV virus (NY-99) diluted in Eegles’s minimal essential medium (MOI, 0.1). A bar graph showing the effect of GZ-161 and GZ-346 on viral release. 24 hours post infection viral release to the media was measured by real-time PCR. Data are means of three replicates ± SEM. *** p <0.001 *versus* infected untreated.

### Antiviral activity of GCS inhibitors against Influenza virus

Since GCS inhibitors showed antiviral activity towards two neuronopathic viruses, and given the current COVID-19 pandemic, we checked whether GZ-161 and GZ-346 will block also non-neuronopathic viruses. The antiviral activity of GCS inhibitors towards the respiratory RNA virus, influenza, was examined. MDCK cells were incubated with 10 μM GZ-161 or GZ-346 one hour prior to infection with mouse adapted influenza virus A/PR/8/34 (H1N1) (PR8). Supernatant were harvested 8 hpi and analyzed by qPCR for the detection of viral RNA in the culture media (Fig 5A). A ∼90% reduction in viral RNA present in the supernatant was apparent in samples treated with GZ-161 or GZ-346 compared to the vehicle DMSO (untreated). In addition to their ability to inhibit PR8 replication, the ability of GCS inhibitors to reduce the cytopathic effect (CPE) of PR8 infected cell was further examined. MDCK cells were incubated with 10 μM GZ-161 or GZ-346 one hour prior to infection with PR8 (MOI, 0.1) and LDH release to the supernatant was measured at 24 hpi as an indication for cell disintegration. Both GZ-161 and GZ-346 significantly reduced PR8-induced cytotoxicity by 65% and 90% respectively (Fig. 5B).

**Figure 5.**
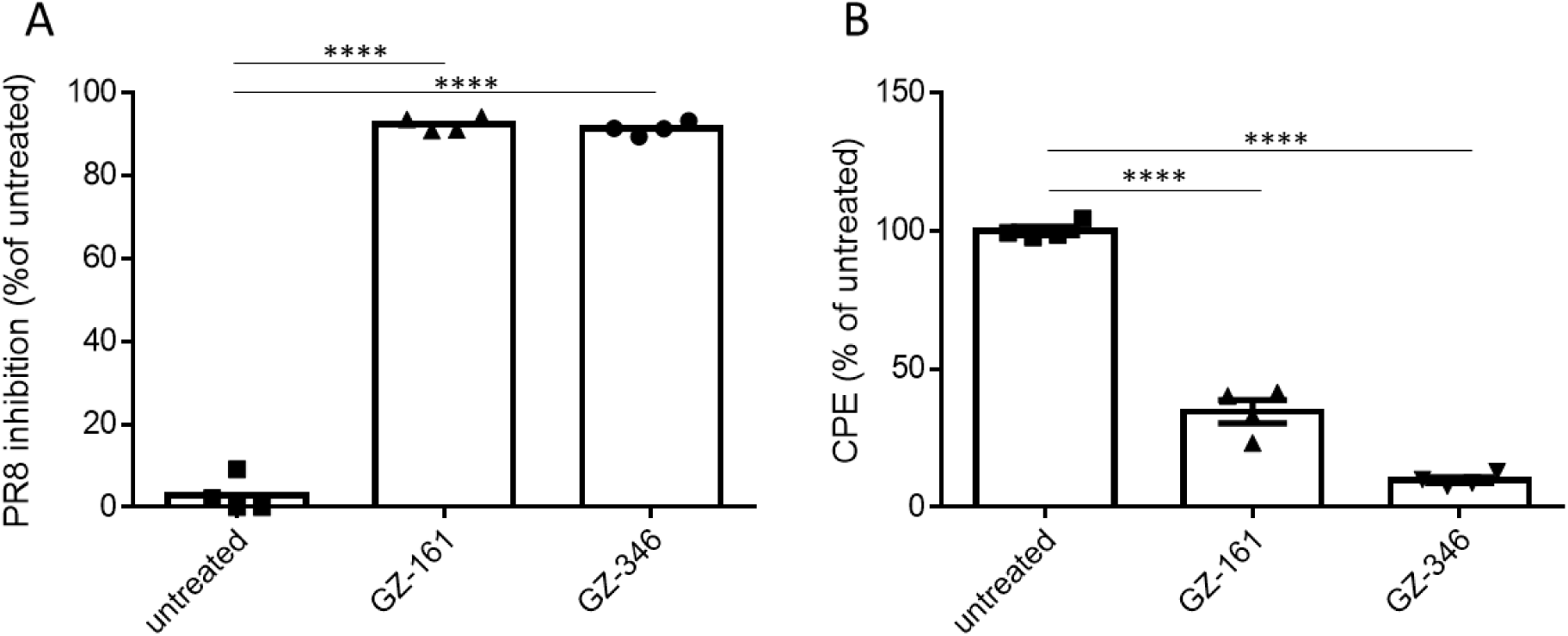
Inhibition of influenza virus A/PR/8/34 (H1N1) by GCS inhibitors. MDCK cells were treated with GZ-161 or with GZ-346 (10µM). 1 hour later, cells were infected with influenza virus A/PR/8/34 (H1N1) diluted in Eegles’s minimal essential medium containing 2 µg/ml trypsin (MOI, 0.1). (**A**) Inhibition of Influenzas virus by glucosylceramide synthase inhibitors. A bar graph showing the effect of GZ-161 and GZ-346 on viral release. Viral release to the media was measured by real-time PCR 8 hours post infection and percentage of inhibition was calculated. Data are means of four replicates ± SEM. (**B**) GCS inhibitors reduce the cytopathic effect of PR8. Cell death was measured 24 hours post infection by LDH cytotoxicity assay kit. Percentage of cytotoxicity were calculated. Data are means of four replicates ± SEM. **** p <0.0001 *versus* infected untreated.

### Antiviral activity of GCS inhibitors against SARS-CoV-2

To test for antiviral activity of GCS inhibitors against SARS-CoV-2, Vero E6 cells were incubated with 10 μM GZ-161 or GZ-346 one hour prior to infection with SARS-CoV-2. Supernatants were harvested 24 hpi and analyzed by plaque forming units assay to measure the effect of the drugs on SARS-CoV-2 replication (Fig 6A and B). ∼ 1.7e7±1.3e6 PFU/ml were detected in the medium of vehicle DMSO (untreated) infected cells, while only 37 ± 23 and 700 ± 339 PFU/ml were detected in GZ-161 and GZ-346 treated cells, respectively, indicating of significant inhibition of virus release (P < 0.0001, P<0.001, respectively). To further determine the efficacy of GCS inhibitors, cells were treated with serial dilutions of GZ-161 or GZ-346 and infected with SARS-CoV-2. Supernatants were collected for qPCR at 24 hpi. As was shown for Vero cells, no toxicity of GZ-161 and GZ-346 was observed at any of the concentrations tested. The IC50 of GZ-161 and GZ-346 treatments were determined at ∼2 μM, under these conditions (Fig 6C). Next, the ability of GCS inhibitors to reduce the CPE of SARS-CoV-2 infected cells in comparison to Remdesivir was examined. Remdesivir, a nucleotide analogue prodrug that inhibits viral RNA polymerases, has shown *in vitro* activity against SARS-CoV-2 (*51*). Due to the current public health emergency, the U.S. Food and Drug Administration (FDA) has issued an Emergency Use Authorization for remdesivir for the treatment of COVID-19 (*52*). Vero E6 cells were incubated with 10 μM GZ-161, GZ-346, or Remdesivir, one hour prior to infection with SARS-CoV-2 and cell viability was measured at 48 hpi. Remarkably, Both GZ-161 and GZ-346 significantly reduced SARS-CoV2-induced CPE. Infection with SARS-CoV-2 reduced cell viability to 40% in the untreated cell, whereas treatment with GZ-161, GZ-346 and Remdesivir increased cell viability to 100%, 100% and ∼75%, respectively (Fig. 6D). Taken together, these results demonstrate that GZ-161 and GZ-346 have an antiviral effect on the SARS-CoV-2 clinical isolate *in vitro*, with a single dose able to significantly inhibit viral replication within 24–48 h.

**Figure 6.**
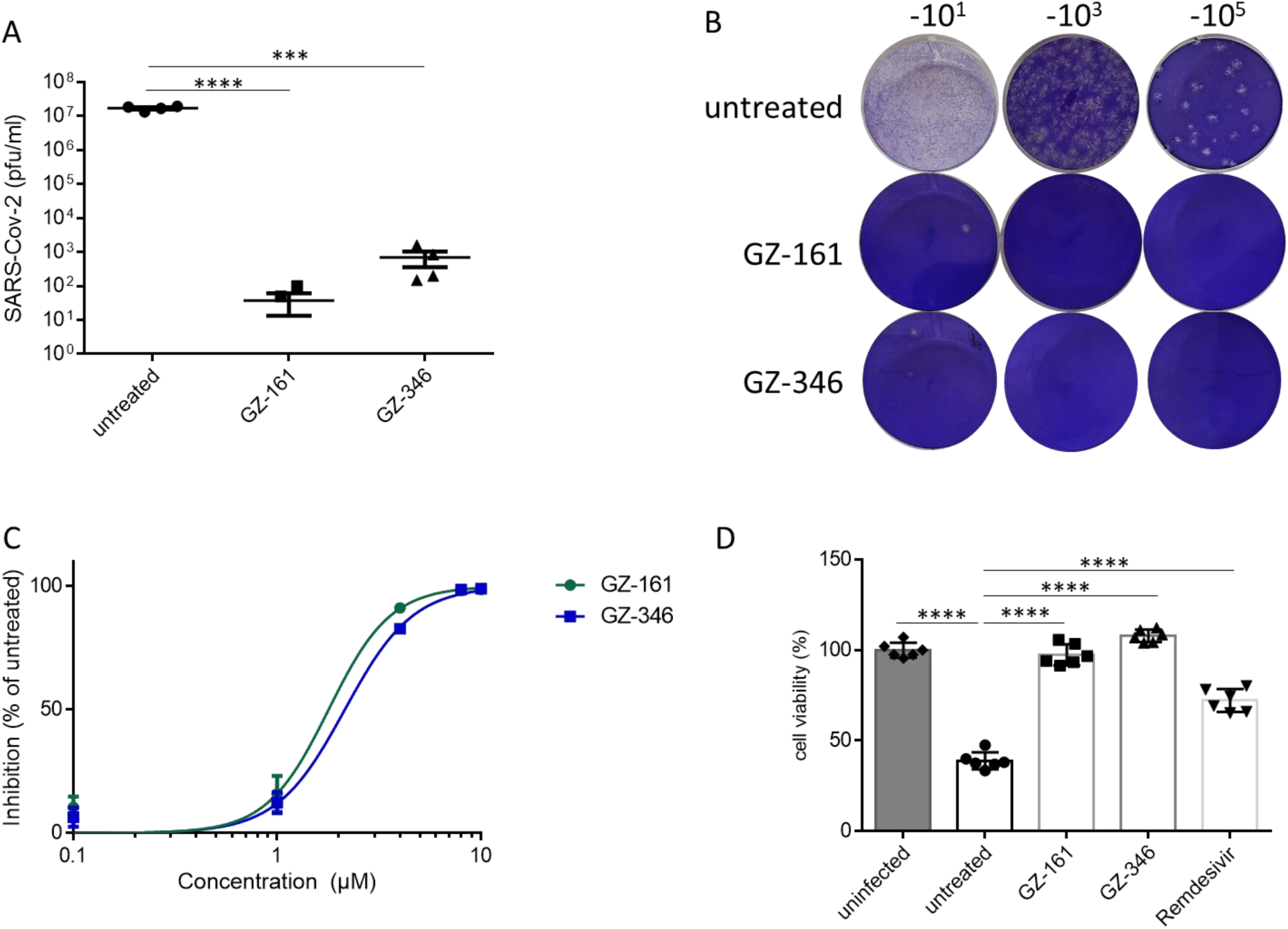
Inhibition of SARS-CoV-2 by GCS inhibitors. **(A, B)** Vero E6 cells were treated with GZ-161 or with GZ-346 (10µM). 1 hour later, cells were infected with SARS-CoV-2 diluted in Dulbecco′s Modified Eagle′s medium (MOI, 0.01). 24 hpi viral release to the media was measured by plaque forming units (PFU) assay. Data are means of four replicates ± SEM. A representative image of PFU assay is presented in (B). (**C**) Dose-response curves of GZ-161 and GZ-346 for inhibition of SARS-CoV-2 infection. Cells were treated with GZ-161 or with GZ-346 (0.1-10µM) 1 hour prior to infection. Viral release to the media was measured 24 hours post infection by real-timeRT-PCR and percentage of inhibition was calculated. Data are means of four replicates ± SEM. A sigmoidal dose–response curve was fitted to the data using Prism GraphPad 6.0 (GraphPad Software). IC50 values are ∼2 µM for both GZ-161 and GZ-346. (**D**) GCS inhibitors reduce the cytopathic effect (CPE) of SARS-CoV-2. 48 hours post infection cell viability was measured by XTT assay kit. Uninfected cells in gray, infected cells in white. Percentage of live cells were calculated. Data are means of six replicates ± SEM. *** p <0.001, **** p <0.0001.

## Discussion

The need for antiviral drugs is real and relevant. This is especially true for diseases without an effective vaccine. Antiviral drugs with a broad spectrum have a particular advantage dealing with emerging disease outbreaks, such as the current COVID-19 pandemic. In this study we demonstrate that the GCS inhibitors GZ-161 and GZ-346 have a broad spectrum antiviral activity. They inhibit *in vitro* viral replication of 4 viruses (SVNI, WNV, Influenza and SARS-CoV-2) from 4 different genus with different routes of infection and target tissues. While in this work we examined the effect of GCS inhibitors against enveloped RNA viruses, their effect on DNA viruses and non-enveloped viruses remains to be elucidated. The mechanism by which GCS inhibitors block viral replication is not fully-resolved. GCS inhibitors inhibit SVNI replication *in-vitro* if administered pre infection, or immediately following it. They prevent GFP expression, which is only expressed after the generation of the replication complex and inhibit the virus also in single-cycle (high MOI) infection. Our data suggest that GCS inhibitors interrupt with early stage of SVNI replication cycle. Sphingolipids play a significant role in endocytosis, thus might play a major role in virus penetration to the cell. Previous works showed that knocking out UGCG (the gene encoding for GCS) impaired the entry of Influenza virus by endocytosis (*43*). In addition to its role in endocytosis, the cellular membrane has a major role in establishing specialized membrane replication compartments (RCs) that are critical sites for the synthesis of the viral genome(*44*). WNV and DENV were shown to stimulate de novo lipid synthesis (*45-47*), while enteroviruses recruit host lipids important for their replication to the RCs by lipolysis (*48*). Whether GCS inhibitors inhibit SVNI cellular entry or the creation of the replication complex remains to be shown. Interestingly, elevation of host Glycosphingolipids, as occurring in the lysosomal storage diseases Gaucher and Krabbe, induces activation of type I interferon response (*6*). Together with our results showing a key role of GlcCer synthesis in the lifecycle of a broad range viruses, it may explain the evolving of antiviral response activation as a result of glycosphingolipids accumulation.

To advance the pre-clinical development of GCS inhibitors as antiviral drugs, we further examined whether GZ-161 is effective *in vivo*. While both GZ-161 and GZ-346 target GCS, only GZ-161 can penetrate the brain, making it the preferred choice for diseases involving the CNS (35). While GZ-161 significantly improved the survival, GZ-346 had no effect when given starting from 2dpi (data not shown), highlighting the necessity of the inhibitor to penetrate the brain in viral-infections of the CNS.

Treatments of SVNI-infected mice with GZ-161 were carried out at 20mg/kg/day, the same dosage that was administered during the preclinical development of GZ-161 for Gaucher disease (*49*) and Fabry (*50*). The preclinical studies using this dosage supported the clinical dossier to approve the safe and effective use of venglustat (GZ-161 analogue) and eliglustat (GZ-346 analogue) in humans at 168 mg/day to treat Gaucher disease. Thus, as a proof of concept, our findings consistently show that GCS inhibitors, at a pragmatic concentration, possesses antiviral activity robust enough to provide protection against severe SVNI infection.

While GZ161 was only tested *in-vivo* on SVNI, *in-vitro* comparison of GZ-161 on the different viruses suggests it is even more potent against Influenza (∼90% inhibition) and SARS-COV-2 (∼100% inhibition) compared to SVNI (∼60% inhibition). Furthermore, the mouse model of SVNI disease is acute with a mortality rate of about 90% and significant CNS damage, and yet GZ-161 treatment was found to be effective.

We therefore believe it is important to test GZ-161 *in-vivo* also against Influenza virus and COVID-19. Naturally, the best means of administration of these drugs for respiratory diseases also needs to be established (Intraperitoneal, intravenous, intranasal, etc.). While treatment with GZ-161 was effective also when administration started post viral exposure, effect was more significant when administration began pre exposure. It is therefore worth considering both therapeutic and prophylactic treatment for populations at high risk.

We show that GCS inhibitors have an antiviral effect on viruses of four different families, suggesting a key role of the glycosphingolipid synthesis pathway in viral infection. Targeting host proteins or pathways utilized by multiple viruses is less prone to the development of resistance to the drug through mutations. It is worth exploring synergies of GCS inhibitors and treatments targeting viral proteins. It is possible that such treatment can lower the required dosage targeting viral protein and lower the risk of viral resistance.

## Materials and Methods

### Cells

Vero (ATCC® CCL-81™), Vero E6 (ATCC® CRL-1586™), Neuro-2a (ATCC® CCL-131™) and Madin-Darby Canine Kidney (MDCK) cells (ATCC® CCL-34™) obtained from the American Type Culture Collection (Summit Pharmaceuticals International, Japan). Cells were used and maintained in Dulbecco’s modified Eagle’s medium (DMEM) supplemented with 10% heat-inactivated fetal calf serum (FCS), Non-essential amino acids (NEAA), 2 mM L-glutamine, 100 units/ml penicillin, 100 μg/ml streptomycin, and 1.25 units/ml nystatin at 37°C under a 5% CO2/95% air atmosphere.

### Viruses

The original strain of Sindbis virus (SV) was isolated in 1990 from mosquitoes in Israel. This strain was used as a source for variants which differ in their neuro-invasiveness and virulence, generated by a serial passages of SV in suckling and weanling mouse brain. At the 15th passage a neurovirulent variant was observed and designated SVN (neurovirulent). After 7 more passages in weanling mouse brains, another variant was observed and designated SVNI (neuroinvasive). The SVNI strain used is both virulent and CNS-invasive [13].

Recombinant Neuroadapted Sindbis virus expressing Luciferase (TRNSV-Luc) was kindly provided by Diane E. Griffin (Johns Hopkins Bloomberg School of Public Health, Maryland).

Recombinant Sindbis virus expressing GFP (SIN-GFP) was kindly provided by Nicolas Ruggli (N. Ruggli and C. M. Rice, unpublished data) (*42*). WNV virus (NY-99, ATCC® VR-1507™) was used.

Influenza virus A/Puerto Rico/8/34 H1N1 (PR8) was kindly provided by Michal Mandelboim (Tel-Aviv University, Israel).

SARS-CoV-2 (GISAID accession EPI_ISL_406862) was kindly provided by Bundeswehr Institute of Microbiology, Munich, Germany. Stocks were prepared by infection of Vero E6 cells for two days when CPE was starting to be visible. Media were collected and clarified by centrifugation prior to being aliquoted for storage at −80°C. Titer of stock was determined by plaque assay on Vero E6 cells monolayers.

### Glucosylceramide synthase (GCS) inhibitors

The compounds GZ-161 ((S)- quinuclidin-3-yl (2-(2-(4-fluorophenyl)thiazol-4-yl)propan-2-yl)carbamate) and GZ-346 ((1R,2R)-nonanoic acid[2-(2′,3′-dihydro-benzo [1,4] dioxin-6′-yl)-2-hydroxy-1-pyrrolidin-1-ylmethyl-ethyl]-amide-l-tartaric acid salt) were obtained from Sanofi. The compounds were stored as 20 mM and 5 mM stock solutions in DMSO or in PBS, respectively, at −20°C until use.

### Inhibition of Sindbis virus in cell-culture

Vero or Neuro-2a cells were seeded at a density of 3 × 10^4^ cells per well in 96-well plates. After incubating overnight, cells were treated in 4-replicates with serial dilution (1-10µM) GZ-161 or GZ-346. Cells were infected 1 hour later with TRNSV-Luc (MOI, 0.01). The infected cells were lysed 23 hours later, and luciferase activity was measured using the Luciferase Assay System (Promega, Madison, WI) by Infinite 200 PRO plate reader (TECAN).

Cell viability of >95% of non-infected cells was determined by XTT assay (Merck, a colorimetric cell proliferation assay, for quantification of cellular proliferation, viability, and cytotoxicity). Evaluation of the half maximal inhibitory concentration (IC50) was performed by GraphPad Prism 6. Percentage of inhibition was calculated by subtracting the ratio of PFU between treated and untreated cells from 1.

### SIN-GFP assays

Vero cells were seeded at a density of 3 × 10^4^ cells per well in 96-well plates. After incubating overnight, cells were treated in sixteen-replicates with 10 µM of GZ-161 or GZ-346. Cells were infected 1 hour later with SIN-GFP (MOI, 5). GFP levels were measured 1, 3, and 5 hours post infection by Infinite 200 PRO plate reader (TECAN).

### Flow cytometry

Vero cells were seeded at a density of 1.5 × 10^5^ cells per well in 6-well plates. After incubating overnight, cells were treated in triplicates with 10 µM of GZ-161 or GZ-346. Cells were infected 1 hour later with SIN-GFP (MOI, 5). 24 hpi cells were collected, stained with Live/Dead cell stain (ThermoFisher, L34955) and live cells were analyzed for GFP signal. Samples were collected using Fortessa Flowcytometer (BD Biosciences) and analyzed with FlowJo software (Treestar).

### Quantitative (real-time) RT-PCR

Supernatant were collected, centrifuged in a table top centrifuge for 5 min at max speed and stored at −80°C. RNA was extracted by qiagen viral rna extraction kit as per the manufacturer’s instructions. RNA load in the media were determined by q RT-PCR. Real-time RT-PCR was conducted with SensiFAST™ Probe Lo-ROX One-Step Kit (Bioline, 78005) and analyzed with the 7500 Real Time PCR System (Applied Biosystems). The PFU Equivalent per ml (PFUE/ml) were calculated from standard curve generated from virus stocks. qRT-PCR primers and probes for the detection of WNV: WN-3NC-F: CAGACCACGCTACGGCG; WN-3NC-R: CTAGGGCCGCGTGGG; Probe WN-3NC: TCTGCGGAGAGTGCAGTCTGCGAT (*53*) qPCR primers and probes for the detection of PR8: PR8-PA-FW: CGGTCCAAATTCCTGCTGA; PR8-PA-RW:CATTGGGTTCCTTCCATCCA; PR8-PA-Probe: CCAAGTCATGAAGGAGAGGGAATACCGCT. qPCR primers and probes for the detection of SARS-CoV-2 N1, obtained from IDT (2019-nCoV CDC EUA Kit, cat #10006606).

### Inhibition of WNV virus in cell-culture

Vero cells were seeded at a density of 5 × 10^5^ cells per well in 6-well plates. After incubating overnight, cells were treated in 4-replicates GZ-161 or GZ-346. One hour later, cells were infected with WNV at MOI 0.1 for 24 h. Supernatant was collected for qPCR. Percentage of inhibition was calculated by subtracting the ratio of PFU between treated and untreated cells from 1.

### Inhibition of Influenza virus in cell-culture

MDCK cells were seeded at a density of 5 × 10^5^ cells per well in 6-well plates. After incubating overnight, cells were treated in 4-replicates with GZ-161 or GZ-346. One hour later, cells were infected in serum-free medium containing 0.5 μg/mL TPCK-trypsin with PR8 at MOI 0.1. Supernatants were collected 8 hpi for qPCR. Cell cytotoxicity was determined 24 hpi by LDH Assay (Cytotoxicity) (ab65393) according to manufacturer’s protocol. Percentage of inhibition was calculated by subtracting the ratio of PFU between treated and untreated cells from 1.

### Inhibition of SARS-Cov-2 virus in cell-culture

Vero E6 cells were seeded at a density of 1.5 × 10^5^ cells per well in 24-well plates. After incubating overnight, cells were treated in 4-replicates with GZ-161 or GZ-346. Cells were infected 1 hour later with SARS-CoV-2 (MOI, 0.01). Supernatant was collected 24 hpi for qPCR and for plaque-forming units (PFU) quantification. For PFU quantification, Vero E6 cells were seeded at a density of 4 × 10^5^ cells per well in 12-well plates. After incubating overnight, cell monolayers were infected with serial dilution of media and 30‒35 plaque-forming units (PFU) of live virus served as control. After two days of incubation at 37 °C, the inhibitory capacity of GCS inhibitors was assessed by determining the numbers of plaques compared with untreated cells. Cells cytotoxicity were determined 48 hpi by LDH Assay (Cytotoxicity) (ab65393) according to manufacturer’s protocol. Percentage of inhibition was calculated by subtracting the ratio of PFU between treated and untreated cells from 1. All experiments involving SARS-CoV-2 were conducted in a BSL3 facility in accordance with the IIBR regulation.

### Studies in mice

C57BL/6 mice (21-days old) were infected intraperitoneally (i.p) with SVNI (15 PFU/mouse). Beginning at 16 days of age (5 days pre-infection) or at 23-days of age (2 days post-infection), injections of GZ-161 were administered i.p (20 mg/kg/day). The compound GZ-161 was dissolved in 30 mM citrate buffer normal saline, pH=5 as a 1 mg/ml stock solution. Mice were weighed daily throughout the experiment, and dosages were adjusted accordingly. Control mice received a similar volume of injection buffer without the active agent.

### Statistical analysis

Statistical analyses were performed with a two-tailed unpaired t-test, or as indicated in the legends. P values are indicated by asterisks in the figures as follows: *P<0.05, **P<0.01, ***P<0.001, and ****P<0.0001. Differences with a P value of 0.05 or less were considered significant. The exact value of n, representing the number of mice in the each experiment, is indicted in the figure legends. Data for all measurements are expressed as means ± SEM. For mouse survival, Kaplan–Meier survival curves were generated and analyzed for statistical significance with GraphPad Prism 6.0 [Log-rank (Mantel-Cox) test (conservative)].

## Acknowledgments

We thank Diane E. Griffin for kindly providing TRNSV-Luc, Nicolas Ruggli for SIN-GFP, and Michal Mandelboim for Influenza virus A/Puerto Rico/8/34 H1N1 (PR8). We thank Shai Weiss for safety advisory. We thank Pablo Sardi from Sanofi for providing GZ-161 and GZ-346.

## Author contributions

E.B.V. and T.I designed the research; E.B.V, R.A, H.A, H.T, Y.Y.R, B.P, N.E, S.M, N.P, and T.I., performed the experiments. E.B.V. wrote the manuscript. All authors discussed results and commented on the manuscript before submission.

## Data and materials availability

GZ-161 and GZ-346 obtained from Sanofi through an MTA.

